# Development of species-specific real-time PCR assays for the identification of five European *Rhinolophus* bats

**DOI:** 10.64898/2026.04.21.719147

**Authors:** Patrick G. R. Wright, Marina Bollo Palacios, Thomas Kitching, Daniel Hargreaves, Szilárd-Lehel Bücs, Ivana Budinski, Branka Bajić, Csaba Jére, István Csősz, Isabella Campbell Harry, Thomas Etheridge, Fiona Mathews

## Abstract

The detection and monitoring of bat species using non-invasive sampling and molecular techniques has become increasingly popular in recent years. In Europe, these approaches have been applied to identify horseshoe bats of the genus *Rhinolophus*, which includes five species: *R. hipposideros, R. ferrumequinum, R. euryale, R. mehelyi* and *R. blasii*. While species-specific real-time PCR assays exist for *R. ferrumequinum* and *R. hipposideros*, no unified panel of real-time PCR assays currently enables the identification of all five European *Rhinolophus* species from non-invasively collected samples. Here, we developed five species-specific real-time PCR assays, each targeting interspecies nucleotide variation within the mitochondrial cytochrome *b* gene. To enhance single-base discrimination, RNase H-dependent PCR (rhPCR) primers were employed, incorporating cleavable blocked primers that require perfect complementarity for extension. The assays were applied to droppings non-invasively collected from 18 caves and one church in Serbia and Romania. Of the 149 samples analysed, 131 (88%) yielded successful amplification of *Rhinolophus* DNA. Detection probabilities for the three species identified in the field ranged from 0.49 to 0.82. Occupancy estimates varied, with *R. euryale* showing the highest (0.86; UI: 0.69–0.97) and *R. mehelyi* the lowest (0.23; UI: 0.08–0.43). The assays were capable of detecting up to three species concurrently within a single pooled sample (approximately 15 droppings). These assays are especially valuable for detecting *R. mehelyi*, given its rarity and uncertain distribution, and offer a robust tool for monitoring *Rhinolophus* populations across Europe.

## Introduction

Non-invasive genetic sampling is a critical tool in wildlife conservation as it can provide information on a species’ presence, population size or ecology without the need to disturb individuals. The detection and monitoring of bat species through the use of molecular techniques has been popularised over recent years as a result of their high level of protection in certain parts of the world and the difficulties in accurately identifying certain species from acoustic data or up close.

Real-time PCR (or qPCR) and metabarcoding are currently the most commonly used techniques for the non-invasive identification of species. Both techniques can be applied to a multitude of sample types for eDNA studies (e.g. water, air) or the collection of animal matter (e.g. fur, droppings) and can provide reliable information on the presence or absence of elusive species (Harrington et al. 2019; Abbas et al., 2026; Rees et al., 2014; O’Neill et al., 2013). While metabarcoding techniques can provide information on all species present within a collected sample, it is also costly and requires a significant amount of pre- and post-sequencing analyses. On the other hand, when targeting few species, real-time PCR offers significant advantages, such as its high sensitivity to very small quantities of degraded DNA and the lack of post-PCR processing (O’Neill et al., 2013). They also allow the identification of multiple species within a single sample when using a set of species-specific primers. This can be advantageous when surveying for bats as many species exhibit mixed roosting behaviour.

All five of the European *Rhinolophus* species occur in southern Europe. Yet, despite the high number of species, little is known about their distributions as they often use vast amounts of unexplored subterranean habitat, with few biologists able to survey for presence or abundance. The three medium-sized horseshoe bat species (*Rhinolophus blasii, R. euryale* and *R. mehelyi*) are of particular interest as they are conservation priorities (EU habitats directive, EUROBATS). Unlike *R. ferrumequinum* and *R. hipposideros*, which are more widely spread in Europe, these species are almost entirely dependent on limestone cave-rich karst areas, and as a result strongly overlap in range but also often utilise mixed-species roosts making their identification at time particularly challenging.

Errors in the identification of *Rhinolophus* bats at specific sites is likely and can result in spurious population trends and misleading information on a species’ distribution (Puechmaille, 2023; Russo, Cistrone & Waldien, 2025). This is particularly notable for *R. mehelyi*, which is listed as vulnerable by the IUCN Red List globally, and endangered at the European level (Russo & Cistrone, 2023). It is thought to be declining with multiple populations thought to have gone extinct in north-eastern Spain, Mallorca, Israel and France (Puechmaille, 2023). While the decline of the species is not contested in Europe, little evidence is available to confirm such trends and difficulties in the identification of the species means that they can be unreliable. Indeed, *R. mehelyi* is morphologically similar to *R. euryale* and both species can be misidentified especially when observations are made from a distance or during hibernation. In France, for example, the most recent reports of hibernating *R. mehelyi* were suggested to be misidentified *R. euryale* after genetic identification (Puechmaille, 2014). In a study by Puechmaille and Teeling (2014), molecular identification of *R. euryale* and *R. mehelyi* was successful but required post-PCR processing and sequencing of a large fragment of mitochondrial DNA. In contrast, Budinski et al. (2024) developed a PCR-based technique that does not require sequencing and enables differentiation of three medium-sized horseshoe bat species based on species-specific band patterns observed on agarose gel. Although this method is fast and cost-effective, it requires the amplification of large DNA fragments making it unfavourable for the identification of noninvasively collected samples. The aim of this study was (1) to develop a real-time PCR assay for the rapid and reliable identification of all European *Rhinolophus* bats and (2) apply the technique to better inform on the distribution of these species, especially *R. mehelyi*, in Romania and Serbia.

## Methods

### Assay development

#### DNA extraction

Tissue biopsies (2-3 mm wing membrane punches) were taken from live bats and stored in 100% ethanol until DNA extraction. Genomic DNA was extracted with the Quick-DNA Miniprep Plus Kit (Zymo Research, USA) following the manufacturer’s solid-tissue protocol for wing punches. For droppings, 1ml of Genomic Lysis Buffer (Zymo Research) was added to each tube, vortexed and incubated at room temperature for a minimum of 30 min. The sample was then centrifuged for 1 minute (min) at 10,000 x g before taking 650µl of supernatant and proceeding from step 4 of the solid tissue extraction protocol. All DNA extracts were then stored at -20°C.

#### Primer Design

The mitochondrial *cytochrome b* (cyt b) gene has been successfully used for the genetic identification of bat species using real-time PCR primers and thus was chosen as the target for this study (Harrington et al 2019). The cyt b gene for *R. blasii, R*.*euryale* and *R. mehelyi* was PCR amplified from tissue sample DNA obtained in this study using primers Bat05A (5’- CGACTAATGACATGAAAAATCACCGTTG-3’) and Bat14A (5’-TATTCCCTTTGCCGGTTTACAAGACC-3’) (Martins et al. 2007). The PCR reaction was performed at a final reaction volume of 20 µl consisting of; 10 µl 2x SYBR qPCR Lo-ROX Mix BLUE (Apto-Gen), 1 µl Bat05A/Bat14A primer mix (2 µM each), 1 µl DNA and 8 µl H_2_O. PCR product formation was monitored on an AriaMX real-time PCR machine (Agilent Technologies) using the following cycling conditions: 94°C for 3 min, followed by 35 cycles of denaturation at 94°C for 45 seconds (s), annealing at 53°C for 45 s, and extension at 72°C for 1 min 30 s, with a final extension at 72°C for 10 min. PCR products were purified using a NucleoSpin Gel and PCR Clean-up kit (Macherey-Nagel) and submitted for external Sanger sequencing (Source BioScience, UK). Raw sequence chromatograms were trimmed and assembled in SnapGene software (Dotmatics, USA).

Tissue derived cyt b sequences along with publicly available sequences retrieved from GenBank were aligned in SnapGene using the Clustal Omega algorithm to generate a consensus sequence for each species. For *R. ferrumequinum* and *R. hipposideros*, consensus sequences were generated exclusively from GenBank data. GenBank sequences were selected from individuals sampled as close as possible to the survey regions. A multiple-sequence alignment of the five consensus sequences was then performed to identify species-specific single nucleotide polymorphisms (SNPs) for use in primer design.

PCR primer pairs for each species were manually designed, incorporating deliberate mismatches to the non-target species near the 31 termini to maximise discrimination. To further enhance assay specificity, RNase H-dependent PCR (rhPCR) technology was employed, with one primer of each pair synthesised as a GEN1 rhPCR primer (Integrated DNA Technologies, USA). The corresponding partner primers were conventional DNA oligonucleotides, designed with additional strategic mismatches to minimise non-specific amplification (Table 1).

**Table 1.**
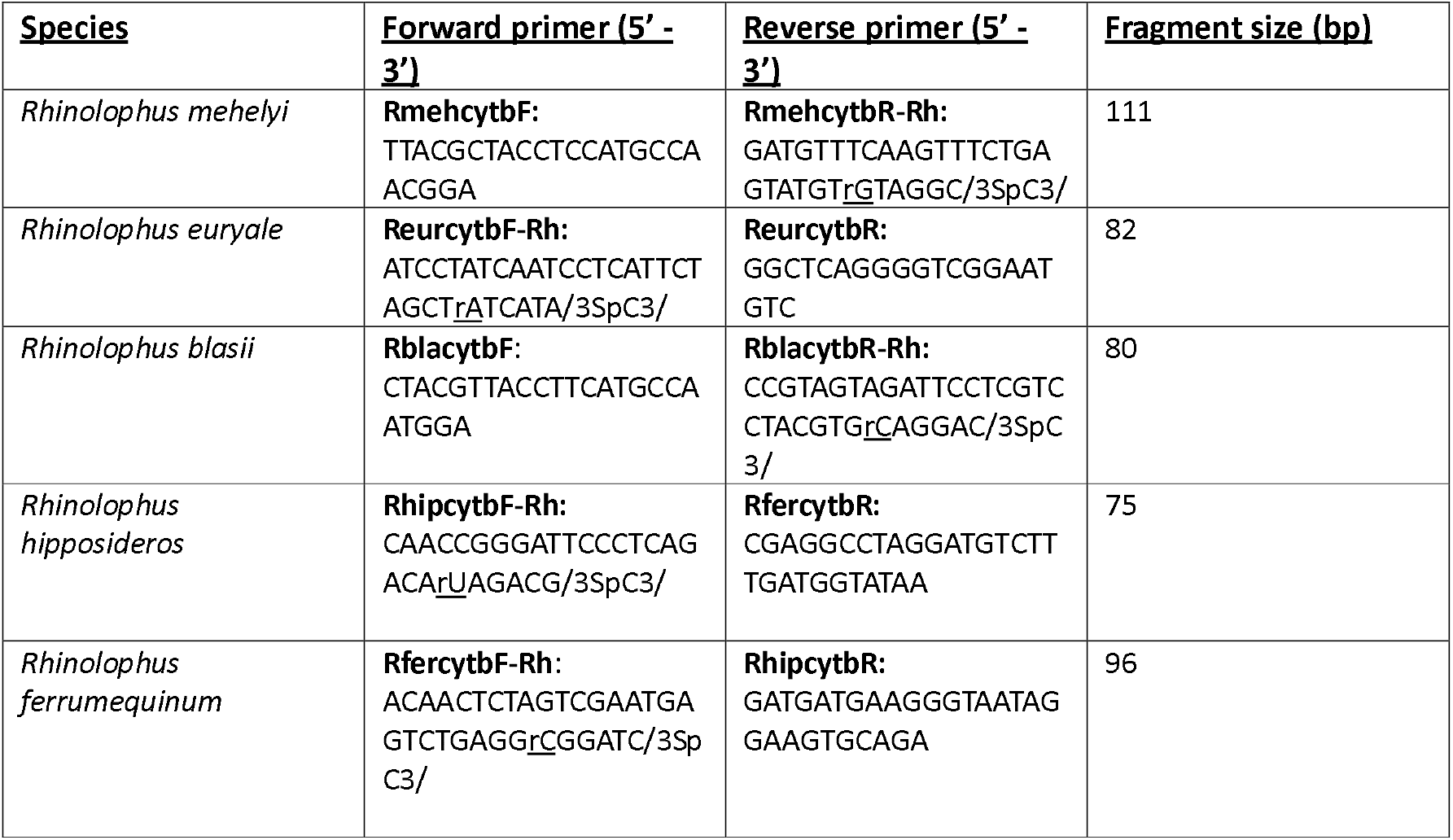
List of species-specific primers designed.

Primer candidates were screened *in silico* with OligoAnalyzer Tool (Integrated DNA Technologies, USA). Where possible, primers were selected to approach the following parameters: a melting temperature (Tm) of 58-62 °C, a GC content 35-65% and ΔG > −9 kcal mol^−1^ for self- or hetero-dimer formation (see Supplementary Table 1)

The specificity of each primer pair was assessed *in silico* using NCBI Primer-BLAST. Each pair was queried against the nucleotide collection (nt) database limited to Chiroptera. Primer pairs were retained only if no significant off-target amplification was predicted for sympatric species.

#### Assay efficiency and specificity testing

To confirm amplification of target sequences and enzyme-dependent activation of the RhPCR primers real-time PCR reactions were conducted against tissue-derived target DNA, with and without the addition of RNase H2. Each PCR reaction consisted of 1 µl DNA, 5 µl 2x SYBR qPCR Lo-ROX Mix BLUE (Apto-Gen, UK), 1 µl primer mix (2 µM forward and reverse), +/- 1 µl 1 mU/µl *P. abyssi* Rnase H2 (Integrated DNA Technologies, IDT) and H_2_O up to 10 µl. PCR amplification was performed on an AriaMX real-time PCR machine (Agilent Technologies) using the following cycling conditions: 95 °C for 1 min, followed by 40 cycles of denaturation at 95 °C for 15 s and extension at 62 °C for 30 s. Each reaction was performed in duplicate alongside no-template controls (NTCs).

To assess amplification efficiency and formation of a single PCR product of each primer pair, 5-fold serial dilutions of tissue-derived DNA were amplified in duplicate for each species. Each PCR reaction consisted of 2 µl DNA, 10 µl 2x SYBR qPCR Lo-ROX Mix BLUE (Apto-Gen, UK), 2 µl primer mix (2 µM forward and reverse), 2 µl 1mU/µl *P. abyssi* Rnase H2 (Integrated DNA Technologies, USA) and 4 µl H_2_O.

The resulting qPCR Cq values from these dilutions were used to generate a standard curve, plotting Cq values against the logarithm of relative DNA concentration (Supplementary Figure 1). The standard curve was used to estimate the coefficient of determination (R^2^) and the slope of the curve, which reflects amplification efficiency. Primer efficiency was calculated using the equation:

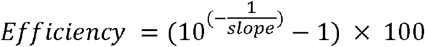

Slopes corresponding to an efficiency range of 90–110%, were considered optimal.

To confirm specific amplification by each primer pair, melt curve analysis was performed with final dissociation step of: 95 °C 20 s, 65 °C for 20 s before gradually heating to 95 °C with a resolution of 0.5°C and soak time of 5 s. The decrease in SYBR Green fluorescence was measured as double-stranded DNA dissociated and the resulting dissociation curve was used to verify the melting temperature of the target amplicon.

To ensure species specificity, each primer set was tested against tissue-derived DNA samples from all five *Rhinolophus* species (Supplementary Table 2). For *R. ferrumequinum* and *R. hipposideros*, tissue samples were provided by Prof. F. Mathews (University of Sussex) from UK specimens. Duplicate PCR reactions were set up consisting of 1 µl DNA, 5 µl 2 x SYBR qPCR Lo-ROX Mix BLUE (Apto-Gen, UK), 1 µl primer mix (2 µM forward and reverse), +/- 1 µl 1mU/µl *P. abyssi* Rnase H2 (Integrated DNA Technologies, US) and H_2_O up to 10 µl. PCR amplification was performed on an AriaMX real-time PCR machine (Agilent Technologies) using the following cycling conditions: 95 °C for 1 min, followed by 40 cycles of denaturation at 95 °C for 15 s and extension at 62 °C for 30 s. For every PCR reaction in this study, primer mixes (2 µM forward and reverse) were made fresh each day from 100 µM stocks (Tris-EDTA pH 8.0 - Integrated DNA Technologies, US). Rnase H2 was also diluted from 5 mU/µL freezer stock to 1 mU/µL in Rnase H2 Dilution Buffer (Integrated DNA Technologies, US) before each experiment.

### Field sampling and data analysis

A total of nine caves and one church were visited in Romania (five in winter and five in summer) and nine in Serbia (three in winter, five in summer and one in both seasons) in 2022. We aimed to collect fresh droppings on the ground where medium-sized *Rhinolophus* bats were roosting. We collected eight tubes of droppings per site and each tube contained approximately 15 pellets and silica beads to improve DNA preservation. The droppings were then stored in a freezer until DNA extraction which took place within two to eight months after collection. For each tube, we followed the DNA extraction protocol on pools of pellets from each tube and real-time PCR settings described in the Assay development section to identify the species present in each cave.

For each species, we used a Bayesian statistical analysis approach to estimate the detection probabilities and occupancy estimates of each species using each tube of droppings as the binary detection/non-detection data for each sampled site. We estimated the probability of detecting each species when present throughout the study area using the R (v. 4.3.2; R Core Team 2021) package ubms (Kellner et al. 2021) and STAN software (Carpenter et al. 2017) implemented in R Studio (v. 2023.12.1; R Studio Team 2023) with 5,000 iterations, 5 chains and the default burn-in setting of half the number of iterations.

## Results

### Assay development

The aim of this study was to develop real-time PCR primer pairs capable of distinguishing between the five European *Rhinolophus* species, enabling accurate species detection from field-collected guano samples. Primers were designed to target regions of the cyt b gene containing species-specific nucleotide polymorphisms (SNPs), maximizing mismatches between target and non-target species (see Table 1 for sequences, Supplementary Table 1 for primer properties).

Due to the close genetic similarity, particularly between *Rhinolophus mehelyi* and *Rhinolophus euryale*, RNase H-dependent PCR (rhPCR) technology was employed to enhance specificity and reduce the chances of cross-reactivity. This approach uses cleavable blocked primers incorporating a single RNA residue at a SNP site, which is cleaved by a thermostable RNase H2 only upon perfect complementarity, thus reducing the chances of extension and amplification in the presence of mismatches (Dobosy et al., 2011).

New primer sets for *R. ferrumequinum* and *R. hipposideros* were designed despite previously published primers existing (Harrington et al., 2019) to enable all five European *Rhinolophus* species to be identified using a single unified panel sharing common reaction conditions and cycling parameters, simplifying workflow and reducing the potential for inter-assay variability.

*In silico* analysis confirmed that primer sets for *R. mehelyi, R. euryale, R. blasii, R. hipposideros* demonstrated specificity for their respective target species. However, *R. ferrumequinum* primers were shown to also be specific to *Rhinolophus* species *Rhinolophus clivosus*. Further investigation highlighted a 96.4% similarity between cytochrome b sequences for *R. ferrumequinum* and *R. clivosus* (GenBank Accession: OR028845.1). Given that *R. clivosus* is geographically restricted to Africa and the Arabian Peninsula, this cross-reactivity was not considered problematic within the context of this study but should be acknowledged for future research applications.

All primer pairs successfully amplified their target sequences when tested on tissue-derived DNA with PCR reactions conducted in the presence of RNase H2 enzyme exhibiting clear activation of the RhPCR primers compared to controls without enzyme addition (Supplementary Figure 1). No-template controls showed no amplification.

Amplification efficiencies for each primer pair were assessed via amplification from five-fold serial dilutions of target species tissue DNA. Standard curves from the resulting Cq values were used to estimate R^2^ values and efficiencies which were all considered optimal, indicating reliable quantitative performance for each assay (Supplementary Figure 2).

Melt curve analysis confirmed the specificity of amplification for each primer pair, with single distinct peaks at consistent melting temperatures corresponding to each target species’ amplicon (Supplementary Figure 2).

Cross-reactivity tests confirmed high specificity of primer sets for *R. euryale, R. blasii, R. hipposideros*, and *R. ferrumequinum*, showing no evidence of non-specific amplification when tested against DNA from non-target European *Rhinolophus* species (Supplementary Table 2). However, the *R. mehelyi* primer pair generated a very late amplification product (Cq 38.42) when tested against *R. ferrumequinum* DNA in both replicates, with a melting temperature (Tm) of 81.5°C. To ensure accurate species identification, positive amplifications for *R. ferrumequinum* were quality-controlled by verifying a Cq value below 34 and a product Tm within the 80–80.5°C range.

### Field test

Of the 149 samples tested, 119 were positive for one or more *Rhinolophus* bat species, while 30 tested negative for *Rhinolophus* DNA. Of these negative samples, 11 produced positive PCR amplification with mammalian 12S primers (Xie et al 2015), indicating the droppings may have originated from non-*Rhinolophus* mammals. Attempts to identify the species from the 12S amplicons were unsuccessful due to poor sequencing quality, possibly resulting from degraded DNA template. A total of 45 samples resulted in the detection of multiple species and 11 detected three species and a single site did not detect any of the targeted species. *R. euryale* was detected in all but two sites and this was illustrated with a very high probability of detection (0.82; UI 0.75-0.88) and occupancy (0.86; UI 0.69-0.97; Figure 1). The probability of detection of *R. mehelyi* (0.61; UI 0.42-0.78) was slightly higher than *R. blasii* (0.49; UI 0.37 0.62), but the probability of occupancy of *R. blasii* was higher (0.43; UI 0.23-0.64) than *R. mehelyi* (0.19; UI 0.06-0.37; Figure 1). The probability of detection of *R. ferrumequinum* was the lowest of all species (0.35; UI 0.22-0.48) and its occupancy was close to *R. blasii* (0.43; UI 0.23-0.65; Figure 1). All three medium-sized horseshoe bats as species were detected within sites along the Romanian and Serbian border in the Iron Gates National Park (Figure 2). *R. mehelyi* was confirmed in a new site in Romania where only *R. euryale* and *R. blasii* were known to be present.

**Figure 1.**
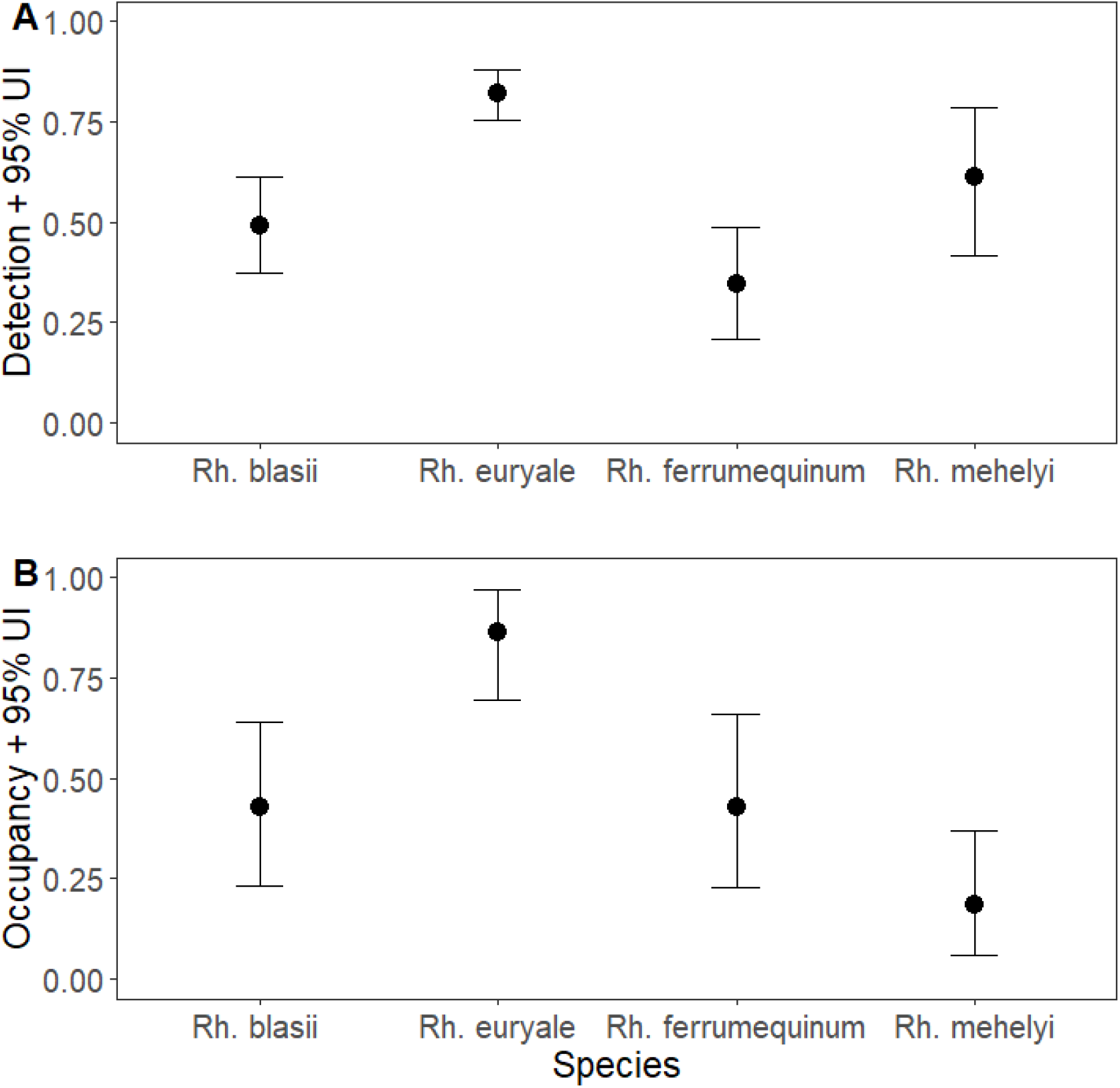
Probability of detection (A) and probability of occupancy (B) with the Bayesian Uncertainty Intervals of all species detected.

**Figure 2.**
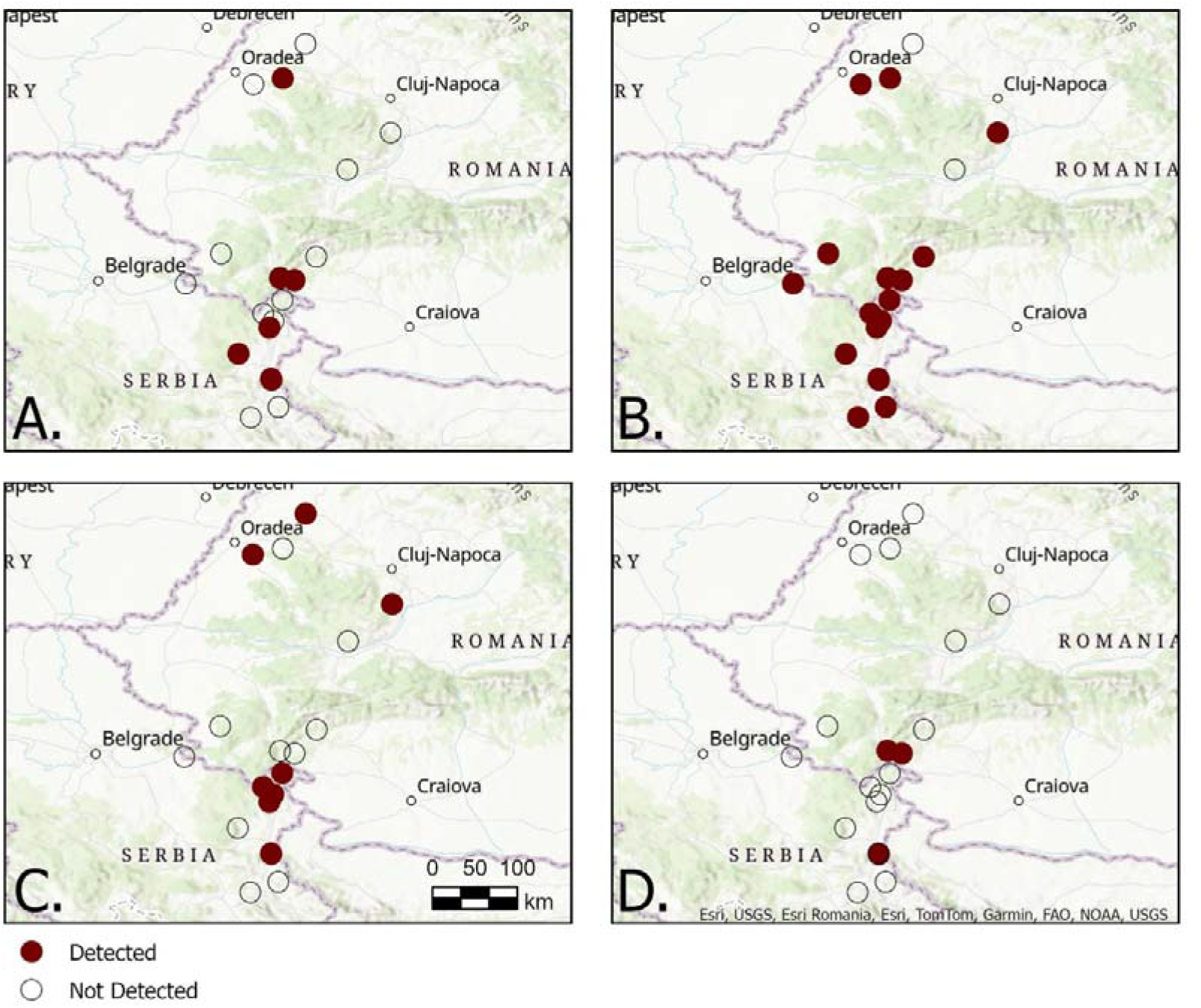
Sites where A. *R. blasii*, B. *R. euryale*, C. *R. ferrumequinum* and D. *R. mehelyi* were detected.

## Discussion

Recent advancements in DNA analysis have significantly enhanced the detection and identification of bat species, enabling more accurate studies of their distributions and ecology. In this study, we developed real-time PCR assays capable of identifying all five European *Rhinolophus* bat species - *Rhinolophus blasii, R. euryale, R. mehelyi, R. ferrumequinum*, and *R. hipposideros* from droppings samples. Field validation demonstrated high amplification success, with few failures, mostly confined to samples collected within a specific time frame, suggesting potential sampling or preservation issues.

Notably, the assay showed a high detection probability for *R. mehelyi* despite its low occupancy across the surveyed sites and confirmed its presence at a new site where its presence was initially unknown. This relatively high detectability is particularly beneficial given the species’ rarity and uncertain distribution, emphasizing the assay’s utility in resolving ambiguities surrounding its presence. We recommend targeted sampling at sites with uncertain *R. mehelyi* records to refine distributional data further.

Although genetic identification of these species can be achieved through direct sampling from individual bats (e.g., wing biopsies, hair samples) using established techniques (Puechmaille and Teeling, 2014; Budinski et al., 2024), the method presented here enables the identification of highly degraded DNA from non-invasively collected samples. Another key advantage of this method is its ability to detect multiple *Rhinolophus* species from pooled dropping pellets, thereby significantly reducing the risk of cross-contamination associated with single-dropping analyses. This method proved effective, with 45 samples detecting multiple species and 11 samples identifying three species concurrently. However, abundant species like *R. euryale* may inadvertently mask the detection of less abundant species due to higher likelihoods of sampling their droppings. Therefore, collecting multiple pooled samples per site is recommended to maximize species detection probability.

Although molecular assays substantially improve bat species identification, there are inherent limitations. Firstly, this assay has not been validated for environmental DNA (eDNA) sampling from substrates such as air or swabs. Further testing focussed on the sensitivity and reliability of the assay would be required to apply it to eDNA studies. Prior to deploying these assays in other regions of Europe, it is recommended that cytochrome *b* sequences are obtained from local populations of each target species to assess whether intraspecific variation at primer binding sites could affect assay performance. Additionally, if deployed in regions with other sympatric *Rhinolophus* species not considered here, further specificity and cross-reactivity tests will be essential. Notably, *in silico* analyses revealed potential cross-reactivity between primers for *R. ferrumequinum* and the African species *Rhinolophus clivosus*. Although geographic isolation currently limits practical concerns, this cross-reactivity must be accounted for in broader geographic applications. It may be possible to overcome this by re-designing the *R. ferrumequinum* primer set or even applying hydrolysis probe technologies, however this was outside the scope of this study.

In conclusion, the assay presented here offers a robust and cost-effective molecular tool for accurately identifying all five *Rhinolophus* species present in Europe from guano samples. With appropriate considerations for the stated limitations, this assay can substantially inform and improve our understanding their distributions.

## Acknowledgments

We are grateful to UNEP/EUROBATS Projects Initiative for funding this project and input from Henry Schofield and Anita Glover in the development of the project.

Authors declare that there are no conflicts of interest in this study.

## Data availability

All data are available in the Supplementary Materials.

## Contributions

PW, MBP, TK, SLB, IB, BB, TE and FM contributed to the study conception and design. Material preparation and data collection were performed by PW, MBP, TK, SLB, IB, BB, CJ, IC. The lab work was undertaken by FM, TE and ICH. The first draft of the manuscript was written by TE, DH and PW.

## Statements and Declarations

Capturing and sampling of bats for the collection of tissue samples were carried out under the permits issued by the responsible authorities and in accordance with the species-specific recommendations of the Canadian Council on Animal Care. All bats were successfully released in good condition at the capture site immediately after processing.

## Funding

This project was funded by the UNEP/EUROBATS Projects Initiative.

## Notes

### Competing Interest Statement

The authors have declared no competing interest.

