## Supplementary Materials for "Development of species-specific real-time PCR assays for the identification of five European *Rhinolophus* bats"

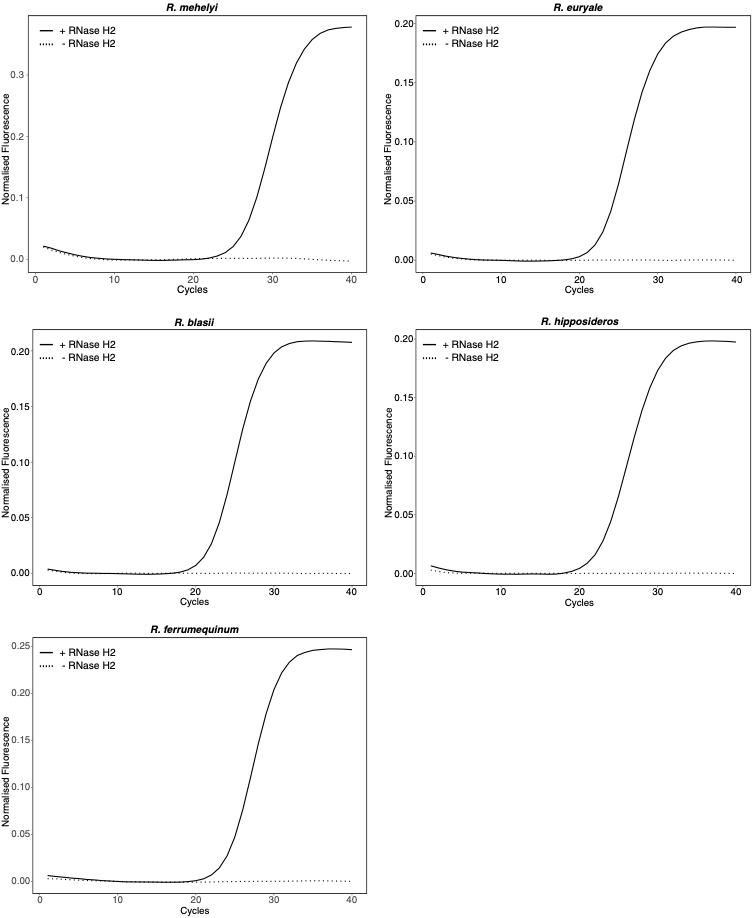

**Supplementary Figure 1** - RNase H2-dependent activation of RhPCR primers for all

five European horseshoe-bat assays.

Real-time PCR reactions were carried out with the species-specific RhPCR primer pair for each target (R. mehelyi, R. euryale, R. blasii, R. hipposideros and R. ferrumequinum). Amplification plots display the accumulation of normalised fluorescence over 40 cycles for reactions performed with RNase H2 (solid line) or without the enzyme (dotted line). In every case, robust amplification is observed only when RNase H2 is present, while reactions lacking the enzyme remain at baseline, confirming that the 3′-blocked RhPCR primers are effectively released by RNase H2 cleavage and that background amplification is negligible in its absence.

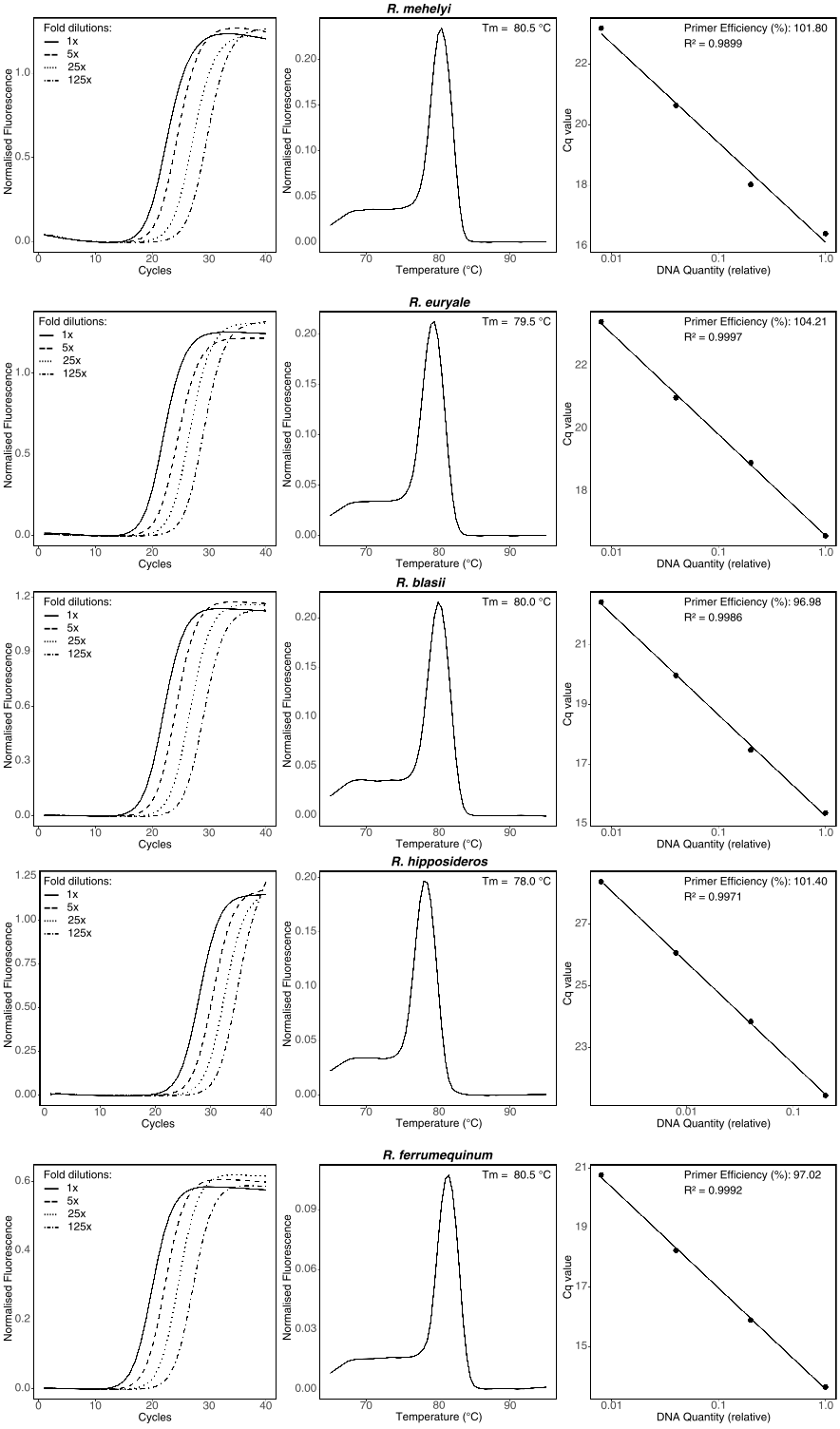

**Supplementary Figure 2** - *Assay development of the five species-specific real-time*

*PCR assays developed for European horseshoe bats.*

Each horizontal row summarises the assay developed for one species. **Left-hand panels:** (amplification plots) show fluorescence accumulation during 40 PCR cycles for a four-point 5-fold dilution series of genomic DNA (1 ×, 5 ×, 25 × and 125 ×; legend inset). **Centre panels:** (dissociation curves) display the first-derivative melt profile of the amplicons, each characterised by a single sharp peak at the

species-specific melting temperature (Tm indicated in the panel header), confirming the absence of non-specific products or primer-dimers. **Right-hand panels:** (standard curves) plot quantification cycle (Cq) values against the logarithm of the relative DNA quantity used in the dilution series. The slope, coefficient of determination (R²) and calculated amplification efficiency (%) are given for each assay.

**Supplementary Table 1 - Primer properties.**

Characteristics and thermodynamic properties of species-specific primers designed for the real-time PCR assay targeting the mitochondrial cytochrome b gene of five European Rhinolophus species. Primer length (nucleotides), GC content (%), melting temperature (Tm, °C), and potential secondary structures such as hairpins and self- or hetero-dimers (with associated ΔG values, kcal mol⁻¹, if applicable) are shown. Rh indicates primers designed using RNase H-dependent PCR (rhPCR) technology. In silico analysis predicted self-dimer formation for two primers (RhipcytbF-Rh and RhipcytbR); however, no adverse effects on amplification were observed during laboratory testing (data not shown).

| **Primer Name** | **Length** | **GC content** | **Tm** | **Hairpin?** | **Self-dimer?** | **Hetero-dimer?** |
| --- | --- | --- | --- | --- | --- | --- |
| RmehcytbF | 24 | 54 | 62 | N | N | N |
| RmehcytbR-Rh | 25 | 36 | **52** | N | N | N |
| ReurcytbF-Rh | 25 | 36% | **52** | N | N | N |
| ReurcytbR | 20 | 65% | 60 | N | N | N |
| RblacytbF | 25 | 48% | 57 | N | N | N |
| RblacytbR-Rh | 25 | 56% | 61 | N | N | N |
| RhipcytbF-Rh | 22 | 55% | 59 | N | **Y (-9.57 ΔG)** | N |
| RhipcytbR | 29 | 45% | 59 | N | **Y (-12.47 ΔG)** | N |
| RfercytbF-Rh | 27 | 48% | 60 | N | N | N |
| RfercytbR | 28 | 43% | 58 | N | N | N |

**Supplementary Table 2** Cross-amplification matrix of the five RhPCR assays

against reference DNA from each European horseshoe bat.

Cycle-threshold (Ct) values are shown for every combination of primer set (rows) and purified genomic DNA from the five target species (columns). Bold values mark the homologous primer-template pairs that produced robust amplification within 40 cycles.

Non-target templates either failed to amplify (n.a., no detectable fluorescence by cycle 40) or amplified late (Ct ≥ 35).

|  | Reference DNA | | | | |
| --- | --- | --- | --- | --- | --- |
| Primers | ***R.mehelyi*** | ***R. euryale*** | ***R.blasii*** | ***R.hipposideros*** | ***R. ferrumequinum*** |
| Rmehcytb | **21.59** | n.a | n.a | n.a | 38.42 |
| Reurcytb | n.a | **20.49** | n.a | n.a | n.a |
| Rblacytb | n.a | n.a | **20.23** | n.a | n.a |
| Rhipcytb | n.a | n.a | n.a | **20.64** | n.a |
| Rfercytb | n.a | n.a | n.a | n.a | **21.14** |
